# Simultaneous mental updating in physical space and number space

**DOI:** 10.1101/2020.03.10.985424

**Authors:** Alexis D.J. Makin, Robin Baurès, Sylvain Cremoux, Tushar Chauhan

## Abstract

People can track moving objects after they disappear or become occluded. For instance, they can estimate the position of a car after it goes behind a truck. People can also track through feature space after occlusion. For instance, they can estimate the current number on a hidden stopwatch. Previous work has suggest that a common rate control module paces mental updating in both physical space and number-space. We tested this by comparing single and dual task conditions. On every trial, participants observed a moving target-counter travel clockwise round a circular track. In the then disappeared for 2, 4, 6 or 8 seconds, and beep sounded. In the single task blocks, participants either estimated the final position *or* the final number. In the dual task block, they estimated final position *and* final number. Performance was very similar in single and dual task blocks. This suggests people can extrapolate through physical space and number space simultaneously. This is a challenge to the common rate control model, which implies there should be some dual task interference. We conclude that if a common rate controller exists, it is capable of pacing simultaneous mental simulations in different dimensions.

## Introduction

People can estimate the current location of occluded moving objects. This ability is necessary for a number of everyday tasks and manoeuvres, such as when a car on a motorway moves behind a truck, or a football is occluded by other players [1,2]. In early work, Travis and Dodge [3] found that participants could track occluded moving objects with their eyes, and the synergy between pursuit and saccadic eye movements during occlusion is now well understood [4–10]. Others have investigated behavioural judgements about occluded moving objects. Authors refer to this as ‘time to contact estimation’, ‘prediction-motion’ or ‘motion extrapolation’, partly reflecting their methodology and/or theoretical assumptions [11,12]. Early work explored the effects of speed, distance and acceleration parameters on performance [13–17]. Perhaps mindful of scientific funding priorities in the early cold war era, Gottsdanker [18] implied that this prediction-motion research would help anti-aircraft gunners track enemy warplanes flying behind clouds. More recently, studies have focused on the neurocognitive mechanisms that underpin prediction-motion, as well as documenting several additional influences on performance [19–29].

Makin (2017) attempted to synthesise the fragmented prediction-motion literature. In some prediction-motion tasks, participants respond (e.g. by pressing a button) when the moving target reaches the end of the occluder (a ‘production task’). Here participants can use either *clocking or tracking strategies*. If participants use the *clocking strategy*, they estimate time-to contact just before occlusion [perhaps by extracting the optic invariant “tau”, 30–33], and then count down the resulting temporal representation to delay their motor response appropriately [34]. Conversely, if participants use the *tracking strategy*, they attempt to track the moving target across the occluder with either covert visuospatial attention or overt eye movements. They then respond when gaze or spatial attention reaches the end of the occluder [35]. As well switching between clocking and tracking strategies, participants often employ heuristics, such as ‘if the target is moving relatively fast, press early’ [36]. These complications make it difficult to interpret systematic effects on performance. For example, it is difficult to interpret the observed influence of target threat [37] or previously seen velocities [38,39].

People can also perceive ‘motion’ (i.e. change over time) in various feature spaces, such as number space or colour space [40–42]. It is also possible to extrapolate through such non-spatial or abstract feature spaces. For instance, participants could observe a static disk morph from yellow to blue then disappear, and press when the hidden disk is the same colour as the background. Alternatively, participants could observe a visible numerical counter disappear, then press when the hidden counter is at zero [43]. In the number task, participants may be extrapolate along an abstract mental number line [44].

Makin and Bertamini [46] argued that all these tasks involve a dynamic mental simulation of the occluded process. For accurate performance, it is necessary to update the dynamic mental simulation at the right speed (not too fast or too slow). It could be that a common rate control mechanism controls the speed of mental updating in all dimensions (Common Rate Control model, CRC, Figure 1A). Alternatively, local predictive circuitry within each feature map might suffice (Separate Rate Control model, SRC, Figure 1B). Local predictive mechanisms probably exist [47,48], however Makin and Bertamini (2014) and Makin and Chauhan (2014) found that performance was similar in different feature spaces, and thus tentatively argued for the common rate control account. Makin [11] suggested that local predictive mechanisms cover very short occlusions (< 200 ms) but the common rate controller takes over for longer occlusions.

**Figure 1.**
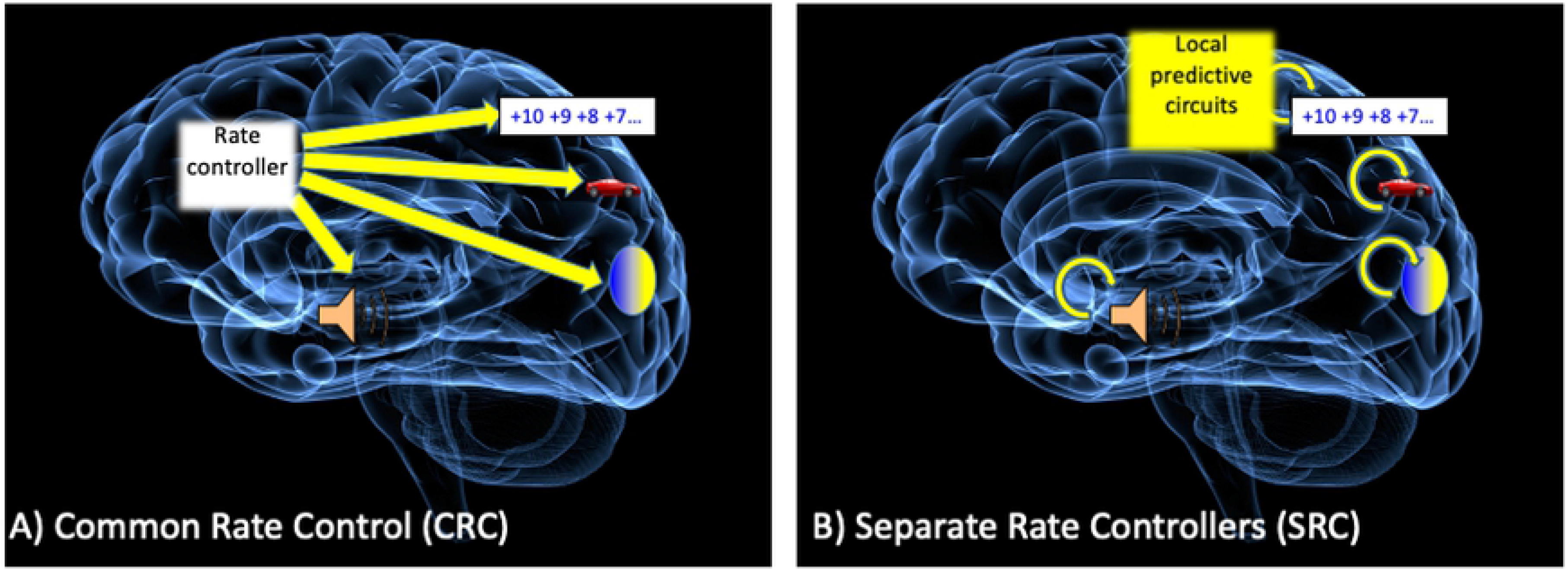
Cartoon versions of the common rate control account (CRC) and separate rate control account (SRC). Adapted from [11].

Despite this previous work, it is unclear whether bottlenecks in the system prevent people from running multiple rate-controlled simulations simultaneously. The common rate control account implies that there should be some dual task interference. This possibility was explored in the current work.

Our tasks are illustrated in Figure 2. A small black number in the middle of a small blue moving target counted up (1, 2, 3, 4…) as it travelled clockwise round a green circular track. On the critical experimental trials, participants observed the visible moving target-counter for 5 seconds. It then disappeared for 2, 4, 6 or 8 seconds. This was followed by a brief auditory tone. Each participant complete three blocks of 80 trials. In one block (Single Position estimation block), they used the mouse to report the estimated final *position* of the hidden target-counter when they heard the beep. Here, they could attend to target position only, and ignore the changing number. In another block (Single Number estimation block), participants used the mouse to report the estimated final *number* on the hidden counter when they heard the beep. Here they could attend to the changing number only, and ignore position. In a third block (Dual estimation block), participants reported the final position *and* number on every trial. Here they had to attend to position and number, and extrapolate through both position and number space simultaneously until they heard the beep. We tested whether performance was impaired in the dual estimation blocks, as the CRC suggests.

**Figure 2.**
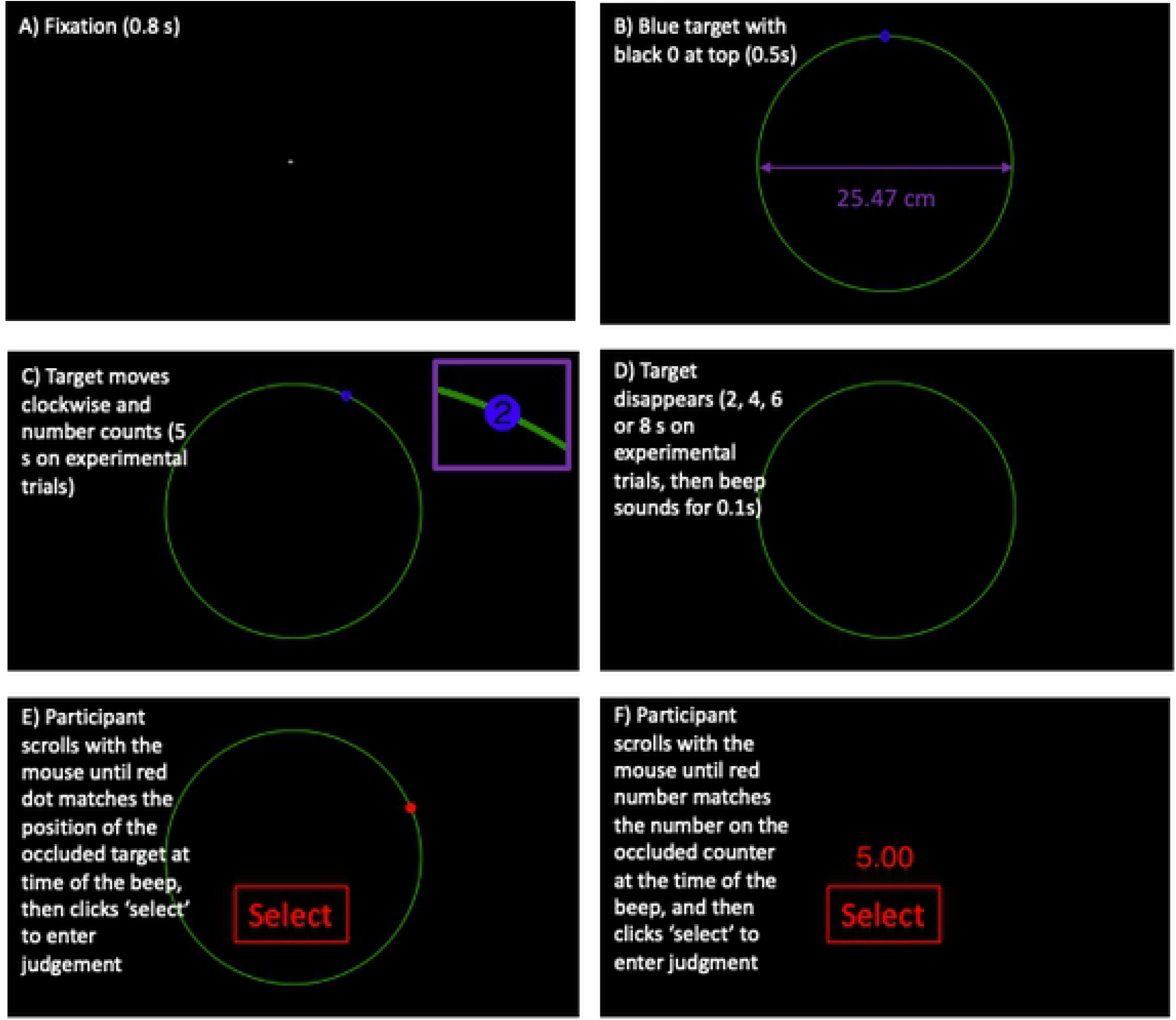
Experimental trial structure. **A)** All trials began with a 0.8 second fixation cross. **B)** This was followed by an 0.5 second interval with an 80 cm circumference (25.47 cm diameter) green circle and the blue target-counter waiting at the start position. **C)** The blue target-counter moved clockwise round the green track, with the counter counting up (see purple inset box for close up). **D)** The target-counter then disappeared for an unpredictable duration and then a 0.1 second beep sounded. Participants then estimated occluded target position at the time of the beep (Single position estimation block, **E**) or occluded number at the time of the beep (Single number estimation block, **F**). In the dual estimation block, they estimated position and then number on every trial (E then F).

## Methods

### Participants

Twenty-four participants were involved in this experiment (age 21 to 32, 2 left-handed, 6 males). All had normal or corrected to normal vision and the experiment was conducted in accordance with the 2008 declaration of Helsinki.

### Procedure

The experiment was programmed in the open-source PsychoPy environment [49] and the code and data have been made available on open science framework https://osf.io/7j5qk/ The stimuli and nature of experimental trials are shown in Figure 2. First a central fixation cross appeared for 0.8 seconds. This was followed by a 0.1 second warning tone (400 Hz) which signalled the start of the trial. The visible target-counter then appeared at the top 12 o’clock position, set a zero, and remained static for 0.5 seconds. It then moved clockwise as the counter incremented in integers (e.g. 1, 2, 3, 4…). Visible motion was followed by a variable occlusion period. The occlusion period was terminated by anther 0.1 second auditory warning tone (800Hz). The visible and occlusion intervals were identical in all blocks.

In the *single position* estimation block, participants used the mouse to enter their estimate of the position of the occluded target when they heard the beep (Figure 2E). A red dot-cursor appeared at occluder onset. Participants scrolled up and down with the mouse wheel to move the red dot cursor clockwise or anti clockwise. When the red dot cursor aligned with their estimate, they clicked the central ‘select’ button. In the *single number* estimation block (Figure 2F) a red number appear centrally (starting at the number at occlusion onset, so ‘5’ in experimental trials). The participants scrolled with the mouse wheel to increase or decrease the red number. When the red number equalled their estimate, they clicked the central ‘select’ button. In the *dual estimation* block, they first reported position, then number. This was because it is relatively easy to hold an auditory representation of estimated final number in memory while scrolling and entering estimated position (e.g. ‘I’m going to enter 11 once I’ve finished with reporting my position estimate’), but relatively difficult to hold an exact visuospatial representation of final occluded position in memory when entering estimated number.

Table 1 shows the spatiotemporal parameters of the *experimental trials,* which were used in all analysis. In experimental trials the visible period was always identical. The blue counter-target travelled 10 cm round the circumference, at 2 cm per second, for 5 seconds. This covered the first 45 degrees of 360 degree circular track. The counter always counted from 0 to 5 in 1 increment per second (the number 5 was not actually seen, because the counter switched from 4 to 5 on the same frame the target became occluded). Occlusion duration (2, 4, 6 or 8 seconds) determined the final occluded angle and final occluded number at beep onset. Note that the final two columns in Table 1 show the correct answers, as given by an ideal observer. Participants were 57 cm from the monitor, so 1 cm = approximately 1 degree of visual angle (to avoid confusion with degrees angle round the circumference circular track, we use cm rather than degrees of visual angle when describing stimulus sizes).

**Table 1.** Spatiotemporal parameters of the Experimental trials. Each block (single position estimation, single number estimation and dual estimation) had 10 of each of these 4 trial types, as well as 40 fillers with randomized parameters (80 trials in total).

| Visible Duration (s) | Visible Distance (cm) | Final Visible Angle | Final Visible Number | Motion speed (cm/s) | Counter Speed (N/s) | Occlusion Duration (s) | Final Occluded Number | Final Occluded Angle |
| --- | --- | --- | --- | --- | --- | --- | --- | --- |
| 5 | 10 | 45 | 5 | 2 | 1 | 2 | 7 | 63 |
| 5 | 10 | 45 | 5 | 2 | 1 | 4 | 9 | 81 |
| 5 | 10 | 45 | 5 | 2 | 1 | 6 | 11 | 99 |
| 5 | 10 | 45 | 5 | 2 | 1 | 8 | 13 | 117 |

Each block also had 40 ‘filler trials’ in order to prevent participants over-learning of these four occlusion durations. On these filler trials, parameters were chosen at random from a bounded uniform distribution, centred on the mean of experimental trials. Target speed was randomized 1-3 cm/s, counter speed was set at 0.5 to 1.5 units/s, visible distance was set at 5-15 cm (30 to 60 degrees round the circumference) and occlusion duration was set at 2 to 8 seconds. Speed, distance and occlusion duration parameters were set independently. The filler trials were not included in analysis.

The single position, single number and dual estimation blocks were completed in a counterbalanced order across the 24 subjects. All were preceded with a short practice block, comprised of 10 filler trials.

### Analysis

First we analysed *estimated final position* and *estimated final number* in single and dual estimation blocks. Given that participants could do the tasks, mean estimated final position strongly correlated with actual final position, and mean estimated final number was strongly correlated with actual final number. Second, we analysed *variable error* (VE). There were 10 trials in each condition, and we computed the standard deviation of estimates across these 10 trials to obtain VE. As VE reflects accumulation of noise across the occlusion interval [50,51] it is observed to increase linearly with occlusion duration [43,46]. Under the CRC model assumptions, we expect VE slopes to be steeper in the dual estimation block, because there is a cost to running two rate controlled simulations simultaneously.

## Results

Results are shown in Figure 3. Performance metrics were very similar in single and dual estimation blocks. As can be seen in Figure 3A and C, performance was similar to that of the ideal observer (green line, intercept = 0 and slope = 1). However, slopes were less than 1, and intercepts greater than zero. This result does not arise from ubiquitous range effects which are often found in prediction-motion tasks (overestimation of low magnitudes, underestimation of high magnitudes [52]). Instead, our participants typically overestimated the final occluded angle (but less so at higher magnitudes), and underestimated the final occluded number (more so at higher magnitudes).

**Figure 3.**
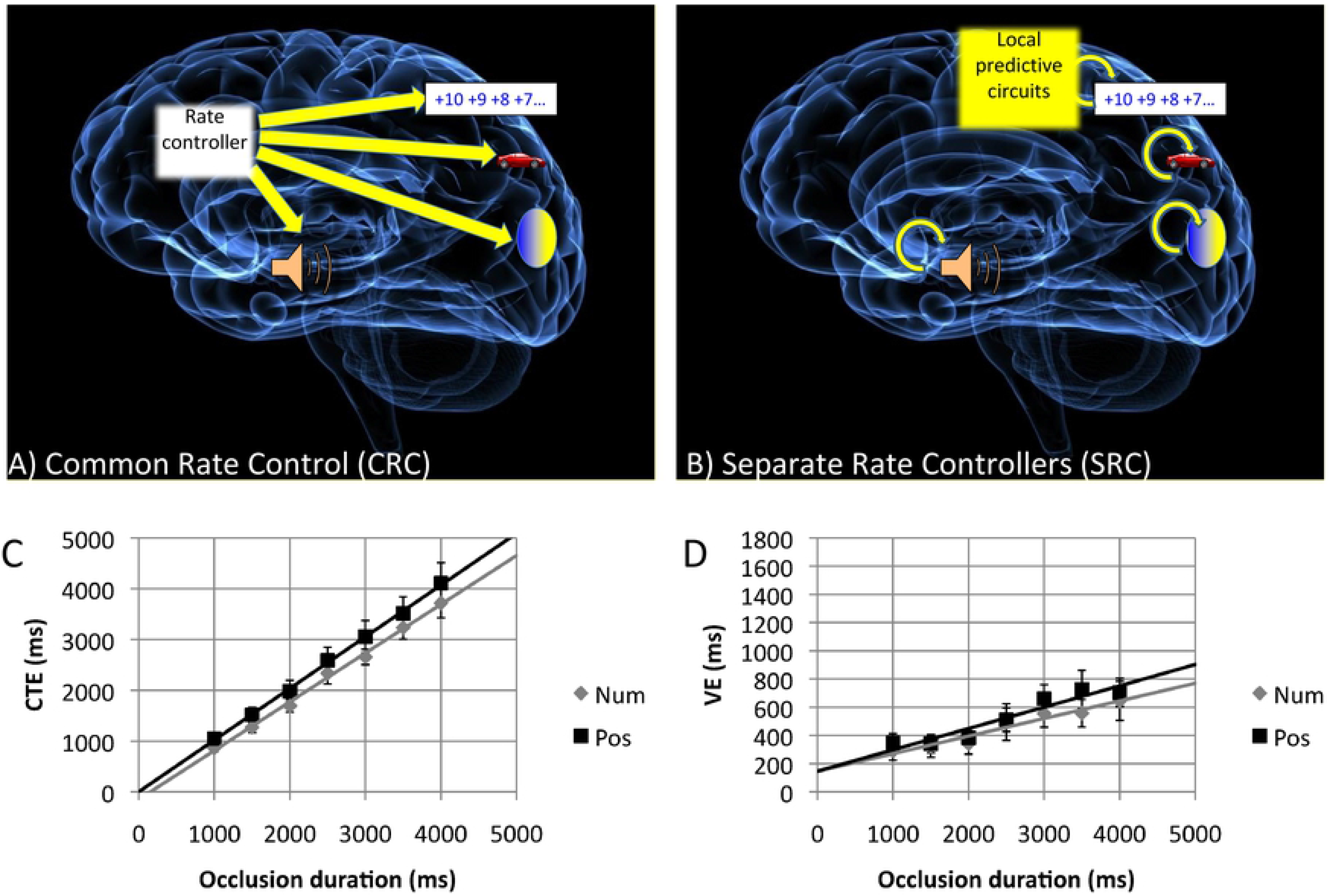
Results and slope analysis. **A)** Final occluded angle vs. Estimated final occluded angle from Position tasks. **B)** Final occluded angle vs. Variable error from Position tasks. **C)** Final occluded number vs. Estimated final occluded number in Number tasks. **D)** Final occluded number vs. Variable error in Number tasks. In all panels red refers to single estimation blocks, and blue refers to results from dual estimation blocks. Regression coefficients are also colour coded. Green lines in A and B show data from ideal observer (i.e. intercept = 0, slope = 1). The four levels of the X axis result from four occlusion durations (2, 4, 6 and 8 seconds). Error bars = +/− 1 S.E.M.

As expected, variable error (VE; the SD of estimates across the 10 repeated trials) also increased with final occluded angle in the position task, and final occluded number in the number task (Figure 3B and D). Final occluded angle and final occluded number were both determined by occlusion duration (2, 4, 6 or 8 seconds, Table 1) in this experiment. Therefore VE increased with occlusion duration, as found in previous work [43,e.g. 50].

### ANOVA analysis

The data shown in the four panels of Figure 3 were analysed with four separate 2 × 4 repeated measures ANOVAs. First, position task data was analysed with 2-factor ANOVAs [2 Block (single, dual) X 4 Final occluded angle (63, 81, 99, 117 degrees)]. Then number task data was analysed with separate 2-factor ANOVAs [2 Block (single, dual) X 4 Final occluded number (7, 9, 11, 13)]. For all ANOVA analysis, the Greenhouse-Geisser correction factor was used when the assumption sphericity was violated.

In the Position task (Figure 3A) there was a large main effect of Final occluded angle on estimated final occluded angle (F (1.208, 27.774) = 234.977, p < 0.001, partial η^2^ = 0.911). There was no difference between single and dual estimate blocks (F < 1), and no Block X Final occluded angle interaction (F < 1).

Again, in the Position task (Figure 3B) there was a main effect of Final occluded angle on VE (F (2.248, 51.708) = 41.947, p < 0.001, partial η^2^ = 0.646), with no other main effects or interactions (largest effect, F (1,23) = 1.119, p = 0.301).

In the Number task (Figure 3C) there was a large main effect of Final occluded number on estimated final occluded number (F (1.369,31.484) = 1292.156, p < 0.001, partial η^2^ = 0.983). However, there was also a main effect of Block (F (1,23) = 19.267, partial η^2^ = 0.456) and Final occluded number X Block interaction (F (3,69) = 3.186, p = 0.029, partial η^2^ = 0.122). The interaction was driven by a growing difference between single and dual blocks with longer occlusions (see slope analysis in next section).

Finally, In the Number task (Figure 4D), there was a large main effect of Final occluded number on VE (Figure 3D F (2.361, 54.308) = 28.595, p < 0.001, partial η^2^ = 0.554). There were no other effects or interactions (largest effect, F (1,23) = 4.000, p = 0.057, partial η^2^ = 0.148).

**Figure 4.**
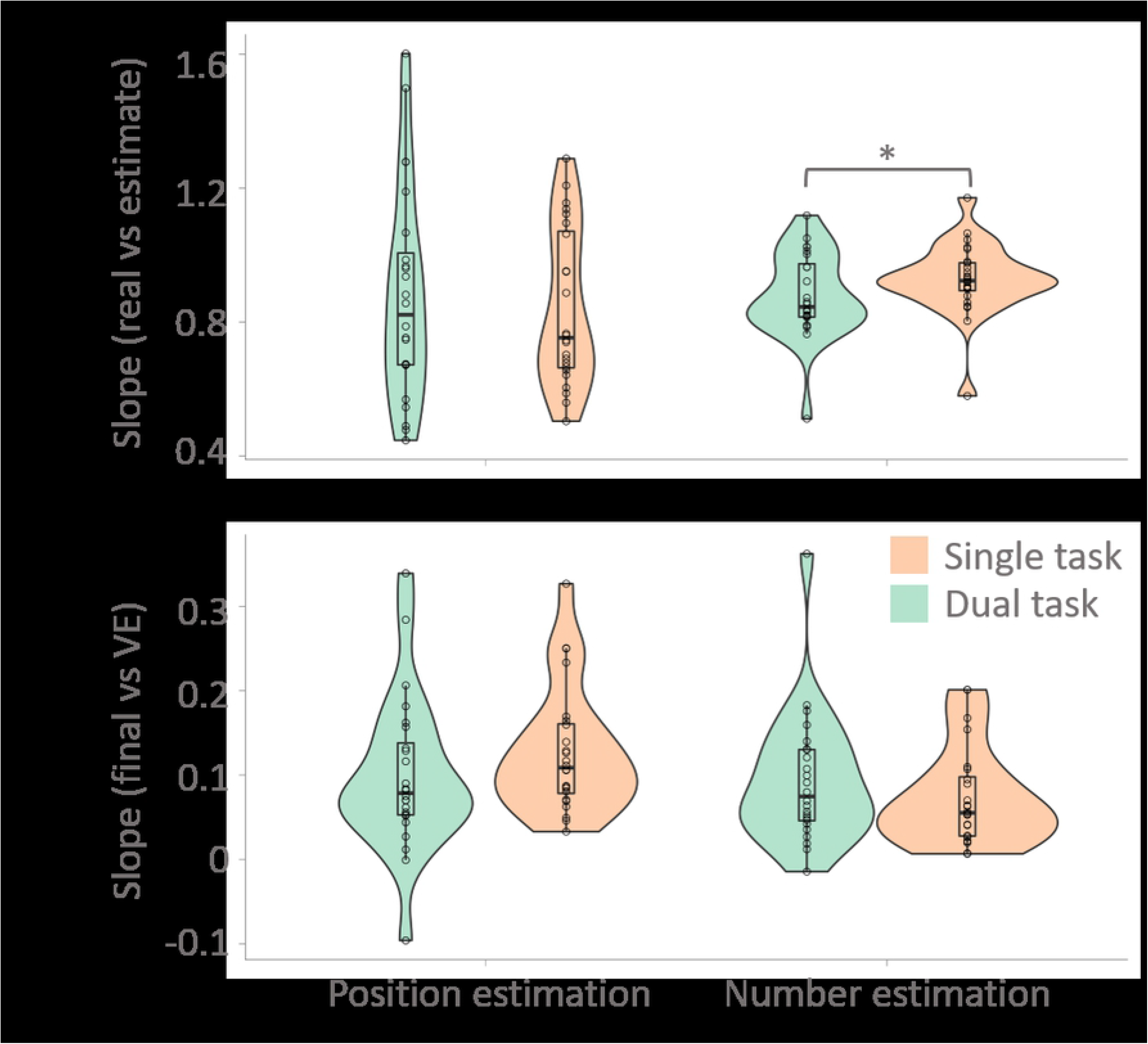
**A)** Slopes of Real final value vs Estimated final value relationships (1 indicates perfect performance). **B)** Slopes of Final value vs VE (0 indicates perfect performance). Dual task is green and single task blocks are orange. Observer data in each condition is shown as small circles, while violin plots show the probability density estimates. Summary statistics are shown as box-plots, with the upper, middle and lower hinge showing the 25th, 50th (median) and 75th percentile. The whiskers extend 1.5 inter-quartile range above/below the upper/lower hinges. The only significant comparison is marked by a * (p < 0.05).

### Regression analysis

To analyse regression coefficients statistically, we extracted intercept and slope values from each participant. Intercepts did not differ significantly between single and dual estimation blocks in any pairwise comparison (largest effect, t (23) 1.586, p = 0.126). Next slope values were analysed with 2-factor ANOVAs [2 Task (Position task, Number task) X 2 Block (single, dual)].

Figure 4A shows the slopes of Final value vs. Estimated final value relationships. There was no difference between single and dual estimation blocks (F < 1, NS), or between position and number tasks (F (1,23) 1.030, p = 0.321). However, there was a Task X Block interaction (F (1,23) = 5.138, p = 0.033, partial η^2^ = 0.183). In the position task, slopes were similar in single and dual blocks (t (23) = 0.800, p = 0.432). Conversely, in the number task, slopes were significantly steeper in the single task block than dual task block (t (23) = 2.645, p = 0.014). We note that this is the only evidence of a dual estimation cost anywhere in our analysis.

Figure 4B shows Final value vs. VE slopes. There were no effects or interactions (F (1,23) = 3.785, p = 0.064). This suggests that VE increased with occlusion duration at approximately the same rate in all conditions.

## Discussion

Participants could simultaneously extrapolate through position and number space with little dual task cost. Underestimation of final occluded number was significantly larger in the dual task block, however the overall pattern of results was one of equivalence. This is a challenge for the common rate control model [11,12] which implies that there should be costly interference effects in the dual task condition. Exactly how challenging is this null result for CRC? If we *had* found robust dual task costs they could be interpreted as confirmatory evidence. However, the lack of interference can be accommodated by the CRC model. It could be that multiple rate-controlled simulations are paced by a single node in the brain (as schematized in Figure 1A). The current results provide evidence which can constrain more detailed versions of the CRC model in future.

We do not think participants used the ‘clocking strategy’ here [35]. There was no way of knowing when the beep would happen, so it was impossible to obtain time-to-contact estimates at occlusion onset. The procedure has many advantages. In particular, it is superior to *production tasks*, where the occluder is visible and participants have to press a button at the end of occlusion interval [22,39]. It may also be superior to interruption paradigms (where participants judge the target reappearance error as being too early or too late) because these can be successfully solved by observers using multiple strategies [11]. We suggest the post-occlusion beep procedure is the best way to force rate-controlled simulation of dynamic occluded processes.

Dual task interference has been demonstrated in other prediction motion studies [53–55]. For instance, the experiments by Baurès, Oberfeld and Hecht [24,56] found that there was a cost to performing two simultaneous prediction-motion tasks. It seems that two simultaneous visuospatial tracking tasks interfere with each other more than one visuospatial tracking and one counting task interfere with each other. When interpreting this, we are mindful that the results of Baurès et al. [24,56] were probably due to dual task effects on subjective velocity *prior* to occlusion. Simultaneous counting did not apparently alter subjective velocity before occlusion in our dual task blocks. Furthermore dual tasking was probably helped by the fact that our counter was on top of the moving target, so there was no need to split spatial attention between task relevant events presented on different parts of the screen.

Although we did not use an eye tracker, it is likely that participants tracked the occluded target-counter in an approximate fashion [8,57]. However, such occluded tracking probably only happened in single position block and dual task block, but not in the number task block. It is known that occluded tracking requires volitional effort after the first 200 ms of occlusion [4,52,58–60], and there was no need to apply this effort in the number task. In future research, these experiments could be replicated with an eye tracker to ascertain whether the dual task requirements have any subtle effect on occluded ocular tracking.

The neural mechanisms which control pursuit eye movements have been investigated humans and monkeys [27,61,62]. Neurons in the extraretinal motion areas MST, IPS and FEFs respond to visible and occluded motion [63–65]. Meanwhile, the DLPFC and the basal ganglia circuitry may be selectively activated during occlusion [7,66]. We know less about the neural mechanisms which guide extrapolation through feature space. The common rate control model predicts that a common node [perhaps in the basal ganglia - SMA “core timing network”, 67], will be activated during occlusion in both position and number tasks [11, Figure 1A]. If the CRC model is broadly correct (despite the current results), then the putative common node may *more* active during dual task conditions, when it is paces two dynamic mental simulations simultaneously. Future neuroimaging studies would be instructive here.

Finally, we note that prediction-motion like tasks could be a useful tool for studying Parkinson’s disease [68], schizophrenia [69,70] or mild traumatic brain injury [71]. It could be that dual task performance is specifically vulnerable to neurological abnormality, even though few dual task costs are found in healthy brains.

